## Supplementary Information for "Predictability and transferability of local biodiversity environment relationships"

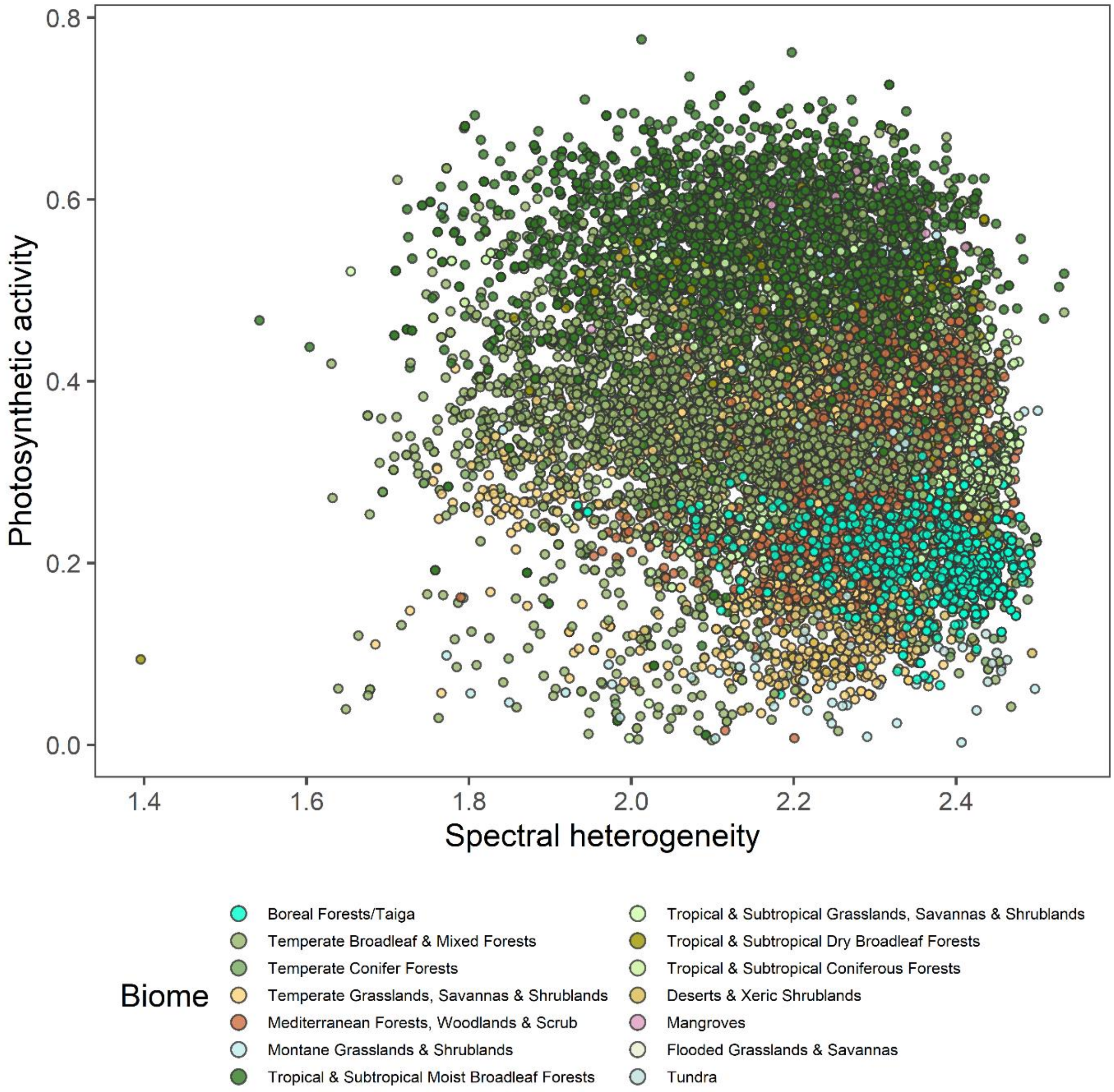


SI Figure 1: Average photosynthetic activity visualized in relation to spectral heterogeneity of each site. Sites coloured by Biome according to Dinerstein et al., (2017).

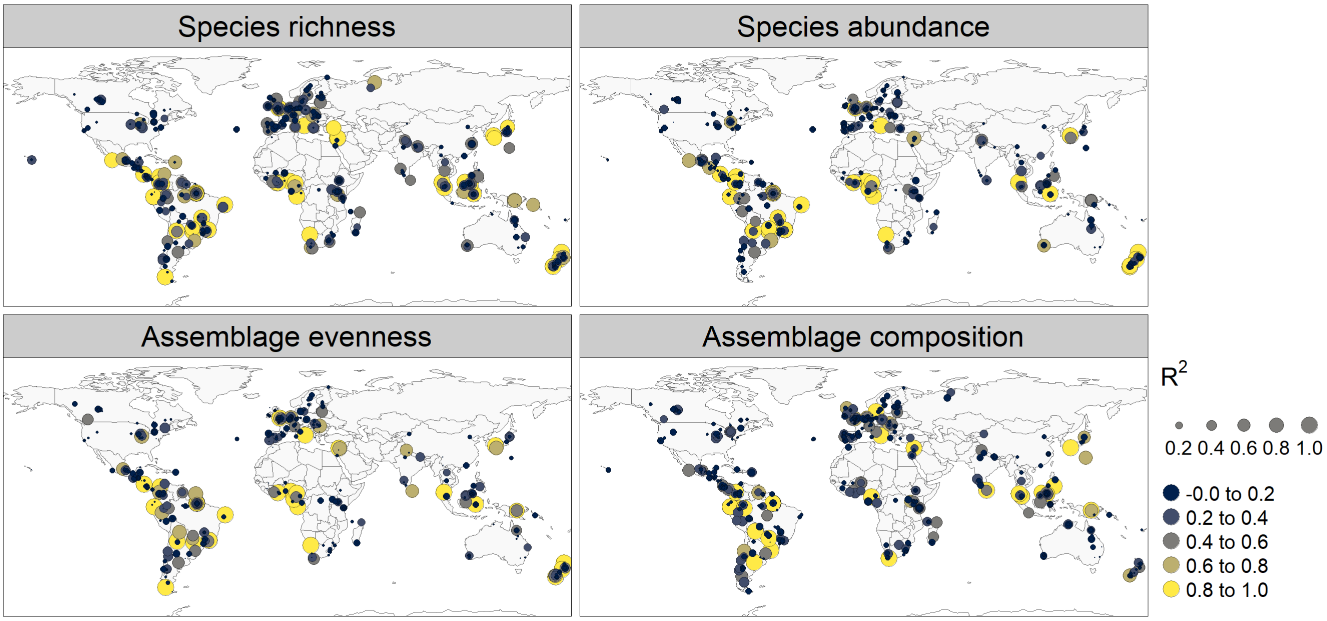


SI Figure 2: Map of average explained variance ($R^2$) of models relating biodiversity measures against environmental predictors. Each dot represents the centre coordinates of a study with size and colour indicating the explained variance.

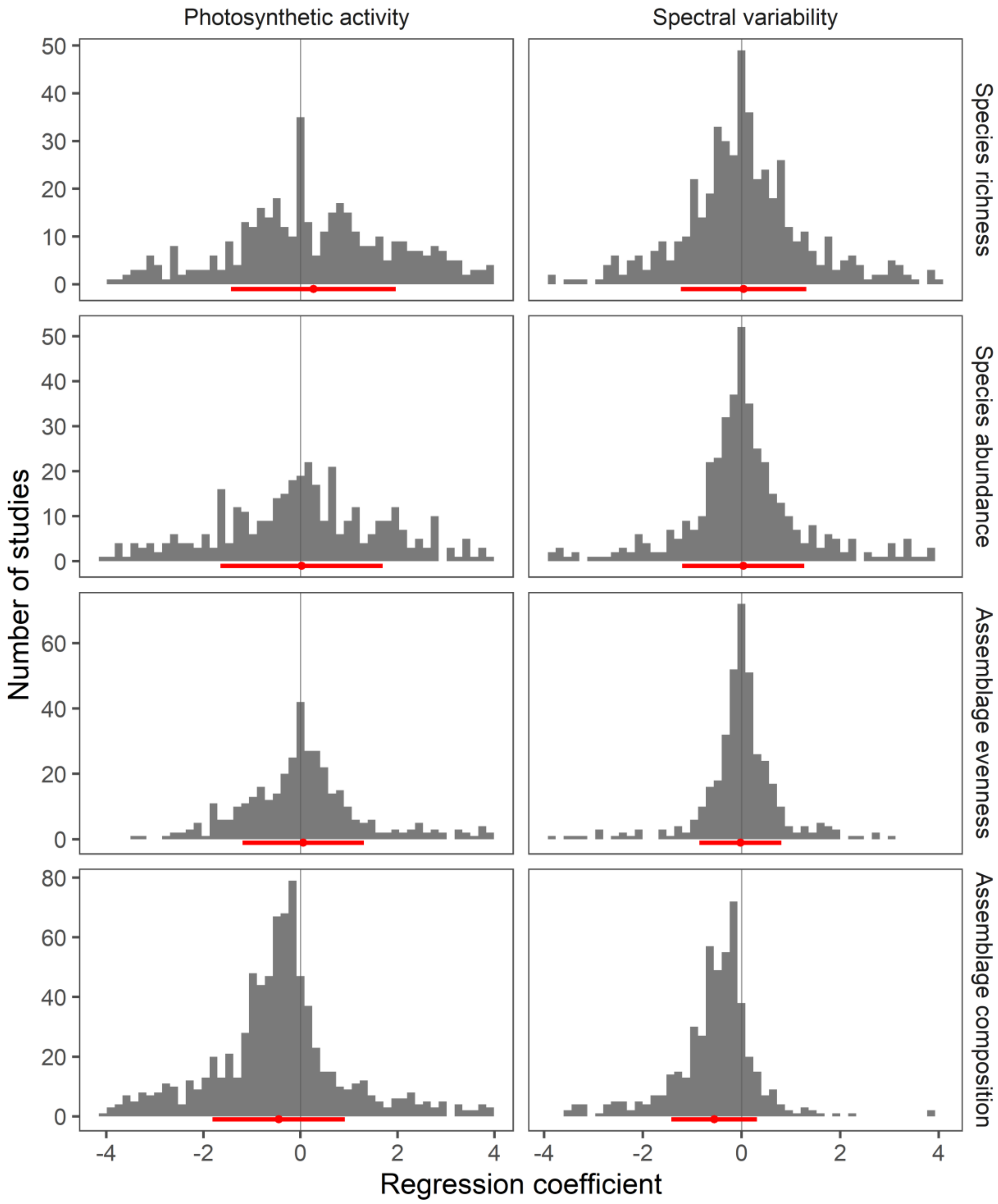


SI Figure 3: Distribution of predictability regression coefficients fitted per study shown for each biodiversity measure (horizontal) and environmental predictor (vertical). Error bars show the mean and 1 standard deviation of the regression coefficients.

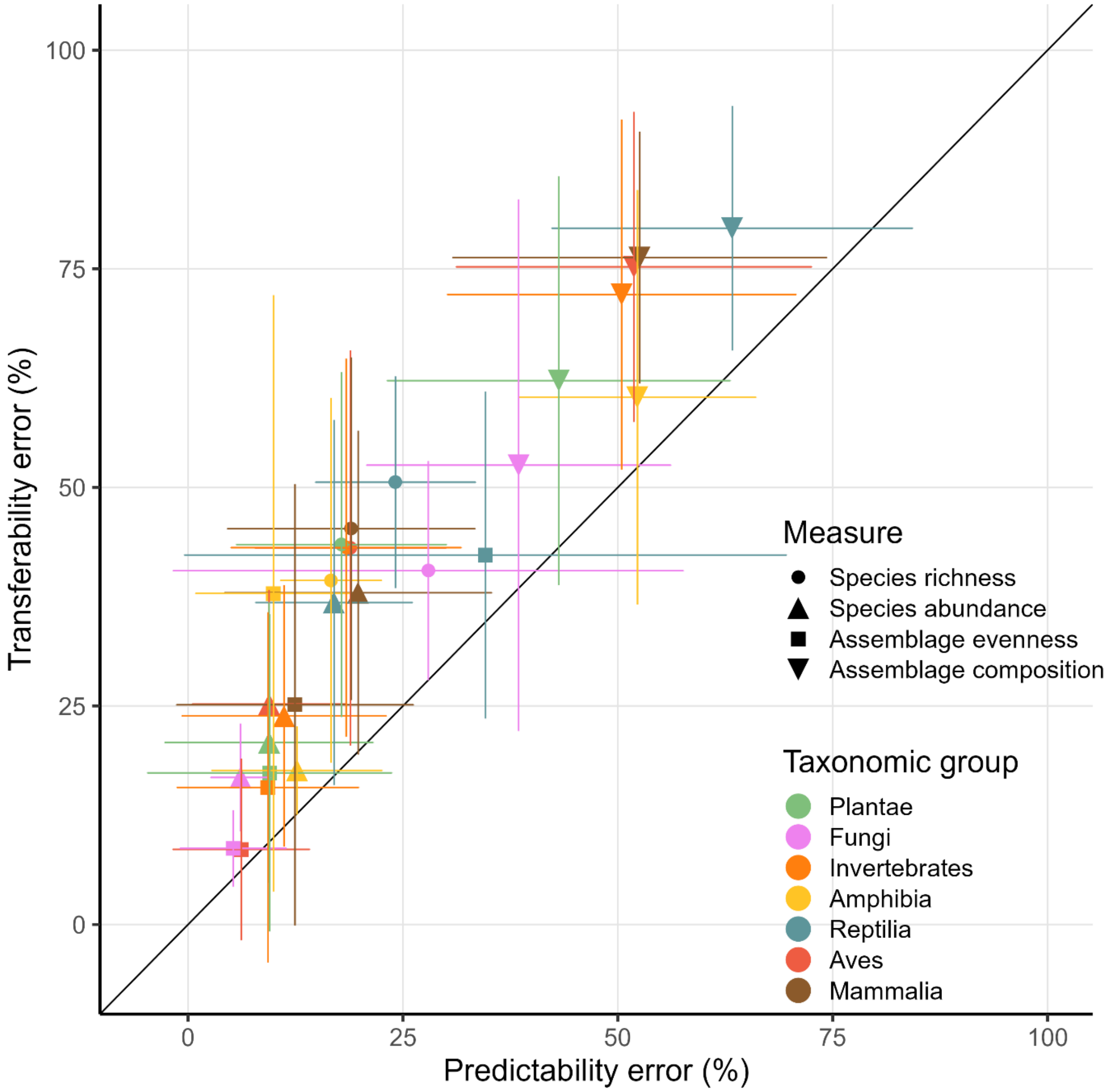


SI Figure 4: Average error (sMAPE) across models for predictability and transferability and using spectral variability as predictor. Colours and shapes as in Figure 4.

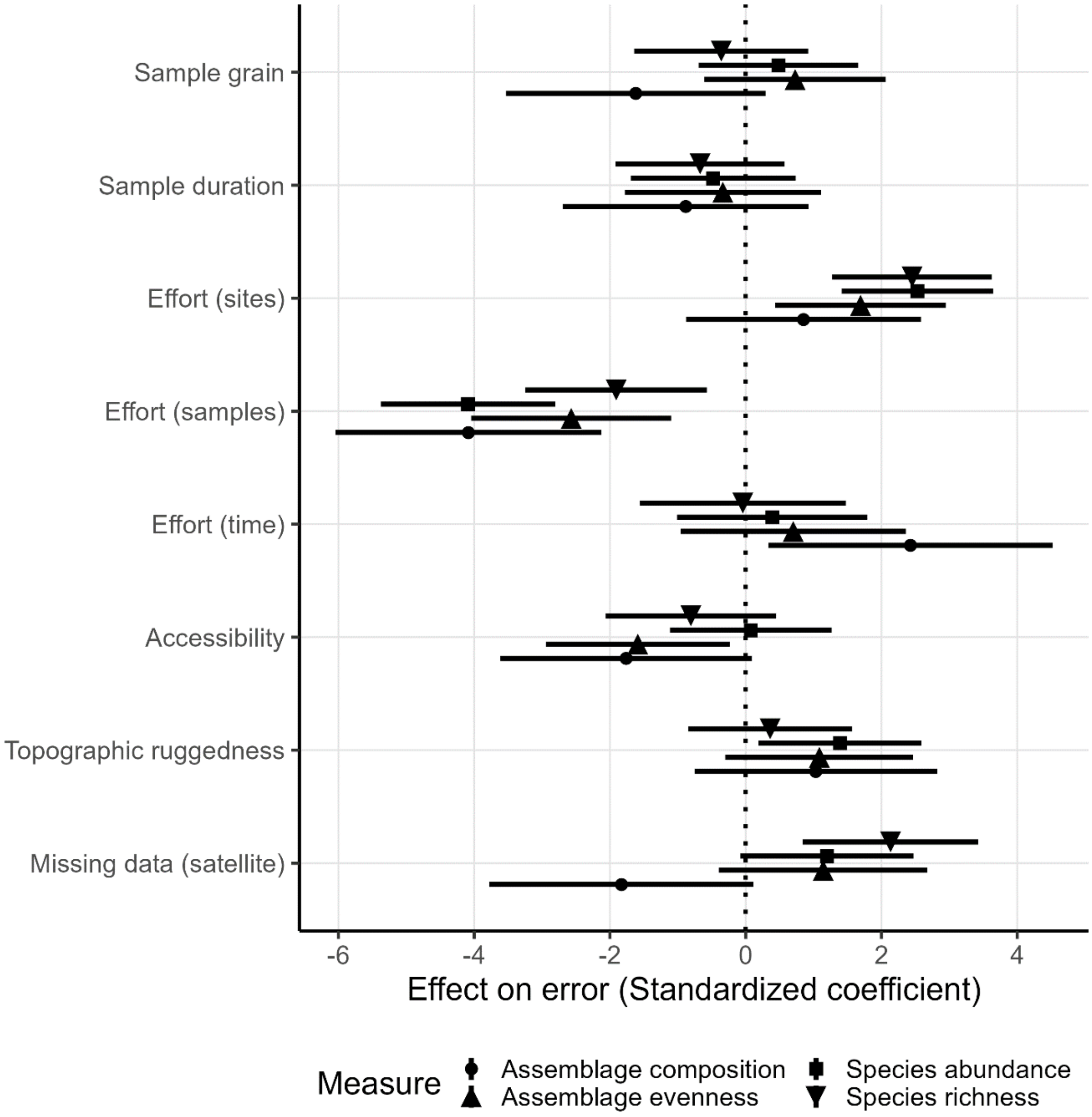


SI Figure 5: Averaged and standardized model coefficients of variables that best explain differences in sMAPE for predictability.

SI Table 1: Combination of sampling methodology pairing used. Taxonomic group indicates the broad taxonomic group used for grouping studies, Sampling method groups together the type of surveying method applied in a study and sampling unit indicates whether effort was measured as number of plots, area covered or time sampled. The concatenation of all columns returns a methodology specific grouping that is used to construct permutations and models.

| **Taxonomic Grouping** | **Sampling method grouping** | **Sampling unit grouping** |
|---|---|---|
| Amphibia | Survey | Time |
| Amphibia | Survey | Plot |
| Aves | FixedPlot | Space |
| Aves | FixedPlot | Time |
| Aves | FixedPlot | Plot |

| | | |
|---|---|---|
| Aves | Transect | Space |
| Aves | Transect | Plot |
| Aves | Survey | Time |
| Aves | Survey | Plot |
| Aves | Trap | Plot |
| Aves | Netting | Time |
| Aves | Netting | Plot |
| Fungi | FixedPlot | Plot |
| Invertebrates | FixedPlot | Space |
| Invertebrates | FixedPlot | Time |
| Invertebrates | FixedPlot | Plot |
| Invertebrates | Transect | Space |
| Invertebrates | Transect | Time |
| Invertebrates | Transect | Plot |
| Invertebrates | Survey | Space |
| Invertebrates | Survey | Time |
| Invertebrates | Survey | Plot |
| Invertebrates | Trap | Time |
| Invertebrates | Trap | Plot |
| Invertebrates | Netting | Time |
| Invertebrates | Netting | Plot |
| Mammalia | FixedPlot | Plot |
| Mammalia | Transect | Space |
| Mammalia | Transect | Plot |
| Mammalia | Survey | Time |
| Mammalia | Survey | Plot |
| Mammalia | Trap | Time |
| Mammalia | Trap | Plot |
| Mammalia | Netting | Time |
| Mammalia | Netting | Plot |
| Plantae | FixedPlot | Space |
| Plantae | FixedPlot | Time |
| Plantae | FixedPlot | Plot |
| Plantae | Transect | Space |
| Plantae | Transect | Time |
| Plantae | Transect | Plot |
| Plantae | Survey | Space |
| Plantae | Survey | Plot |
| Plantae | Trap | Time |
| Reptilia | Survey | Plot |
| Reptilia | Trap | Plot |
